# An inference approach combines spatial and temporal gene expression data to predict gene regulatory networks in Arabidopsis stem cells

**DOI:** 10.1101/140269

**Authors:** Maria Angels de Luis Balaguer, Adam P. Fisher, Natalie M. Clark, Maria Guadalupe Fernandez-Espinosa, Barbara K. Möller, Dolf Weijers, Jan U. Lohmann, Cranos Williams, Oscar Lorenzo, Rosangela Sozzani

**Affiliations:** Plant and Microbial Biology Department, North Carolina State University, Raleigh, NC, USA.; Biomathematics Program, North Carolina State University, Raleigh, NC, USA.; Departamento de Botánica y Fisiología Vegetal, Instituto Hispano-Luso de Investigaciones Agrarias (CIALE), Facultad de Biología, Universidad de Salamanca, C/Río Duero 12, 37185 Salamanca, Spain; Laboratory of Biochemistry, Wageningen University, Dreijenlaan 3, 6703HA, Wageningen, The Netherlands.; Department of Stem Cell Biology, University of Heidelberg, Heidelberg D-69120, Germany; Electrical and Computer Engineering Department, North Carolina State University, Raleigh, NC, USA.

**Author notes:** Present address: Department of Plant Systems Biology, VIB, Technologiepark 927, B-9052 Ghent, Belgium; Department of Plant Biotechnology and Bioinformatics, Ghent University, Technologiepark 927, B-9052 Ghent, Belgium. These authors contributed equally to this work. Corresponding Author: Rosangela Sozzani.

## Abstract

Identifying the transcription factors (TFs) and associated networks involved in stem cell regulation is key for understanding the initiation and growth of plant tissues and organs. Although many TFs have been shown to have a role in the Arabidopsis root stem cells, a comprehensive view of the transcriptional signature of the stem cells is lacking. In this work, we used spatial and temporal transcriptomic data to predict interactions among the genes involved in stem cell regulation. For this, we transcriptionally profiled several stem cell populations and developed a gene regulatory network (GRN) inference algorithm that combines clustering with Dynamic Bayesian Network (DBN) inference. We leveraged the topology of our networks to infer potential key regulators. The results presented in this work show that our combination of molecular biology approaches, computational biology and mathematical modeling was key to identify candidate factors that function in the stem cells. Specifically, through experimental validation and mathematical modeling, we identified *PERIANTHIA (PAN)* as an important molecular regulator of quiescent center (QC) function.

## Introduction

Identifying the transcriptional signature underlying stem cell regulation is key to understanding the initiation and growth of plant tissues and organs. The *Arabidopsis thaliana* root provides a tractable system to study stem cells since they are spatially confined at the tip of the root, in the so-called stem cell niche (SCN), and are anatomically well characterized. The SCN contains several stem cell populations that give rise to the different root tissues and are organized by signals that originate in the quiescent center (QC) (1). Key transcription factors (TFs) in the Arabidopsis root have been shown to be necessary for root formation and stem cell maintenance (2–8). Despite these important findings, a transcriptional signature of the root stem cells is lacking. Genome-wide transcriptional data paired with the development of Gene Regulatory Network (GRN) models can be used to identify additional factors involved in stem cell regulation and predict how genes interact in a molecular pathway. Among the methods to derive dynamic GRNs are Dynamic Bayesian Networks (DBNs), which leverage time series data to infer statistical dependencies among the modeled genes. However, time series datasets alone cannot capture the dynamics of a diverse group of cell populations that are spatially separated, such as the distinct stem cell types. Thus, inferring dynamic GRNs by combining spatial with temporal data becomes necessary for capturing the transcriptional differences between cell types.

To obtain networks of genes that play a role in stem cell regulation, we acquired the transcriptional profiles of several root stem cell types. These profiles allowed us to identify genes that are highly expressed in each of the stem cell populations. We then inferred relationships among those genes to determine their relative importance and their predicted regulatory interactions. To this end, we developed <u>GE</u>ne regulatory <u>N</u>etwork Inference from <u>S</u>patio <u>T</u>emporal data algorithm (GENIST), a DBN-based algorithm capable of integrating transcriptional datasets of different characteristics to reconstruct GRNs. First, we applied GENIST to find groups of genes with similar expression patterns. For this, we used a spatial dataset (QC, CEI (9), XYL, SCN (10)) in combination with transcriptional profiles corresponding to the elongation (Stage II) and differentiation (Stage III) zones of the root (11). Next, we used GENIST to infer regulations among those genes using the transcriptional profile of 12 developmental zones of the Arabidopsis root, which embeds temporal information (12). Finally we represented the inferred GRNs of the stem cell enriched genes using cytoscape (13). The resulting GRNs contained genes that play a role in stem cell regulation and capture their regulations throughout the root ontogenic development. Moreover, our GRN pipeline predicted that a known floral regulator, PERIANTHIA (PAN), is important for QC function. Specifically, phenotypical analyses of a PAN overexpressor and inducible lines, as well as a *pan* mutant, showed that PAN is involved in QC and columella maintenance. Additionally, we obtained the QC specific transcriptional profile in a *pan* mutant to validate our mathematical modeling and to further understand how PAN and its predicted downstream targets are involved in stem cell regulation. Our networks and their analyses are an important step towards defining the genetic framework underlying stem maintenance and function.

## Results

### Identifying root stem cell specific genes from transcriptomic data

To infer GRNs in the stem cells, we first acquired the cell-type specific transcriptional data from the QC cells, cortex/endodermis initials (CEI) (9), xylem initials (XYL) and the whole stem cell niche (SCN) (10)(Methods) (Fig. 1A-D). We then compared these four samples to the transcriptional profiles corresponding to the meristematic (Stage I), elongation (Stage II), and differentiation (Stage III) zones of the root (11) and determined genes highly expressed in stem cells. We found that the expression of many of the known stem cell regulators (SHORTROOT (SHR) (14,15), SCARECROW (SCR) (16,17), MAGPIE (MGP) (18,19), JACKDAW (JKD) (18,19), PHABULOSA (PHB) (5,20,21), PLETHORA 2 (PLT2) (2,22), PLETHORA 3 (PLT3/AIL6) (2,22)) were differentially expressed (Methods) in the stem cells compared to Stage II and Stage III, but were not differentially expressed in the stem cells compared to Stage I. This suggested that important stem cell factors may be expressed in the meristematic zone as well as in the stem cells. We performed a Principal Component Analysis (PCA) to understand if the stem cells and the meristematic cells (Stage I) have similar transcriptional profiles (Fig. 1E). With a cumulative variance of 82.8% in the first 3 principal components, the PCA showed that the variance between the stem cells and the profile captured by Stage I was small. This implied that the differentially expressed genes in the stem cells and Stage I have similar expression patterns. Therefore, to ensure that important regulators were not omitted in our study, we excluded Stage I from our subsequent analysis and identified 1625 genes, containing 201 TFs, as differentially expressed in the stem cell populations versus Stage II and Stage III (Methods) (Fig. 1F) (Dataset S01). Since we observed that regulation of transcription and TF activity were enriched Gene Ontology (GO) categories among these genes (Table S1) we focused our subsequent analyses on the 201 TFs enriched in the stem cells. Among the 201 TFs, 29 had been previously found to have roles in either early embryo development or post-embryonic processes (i.e. stem cell specification), and 16 additional TFs had been reported to be enriched in the stem cells (Dataset S02).

**Figure 1.**
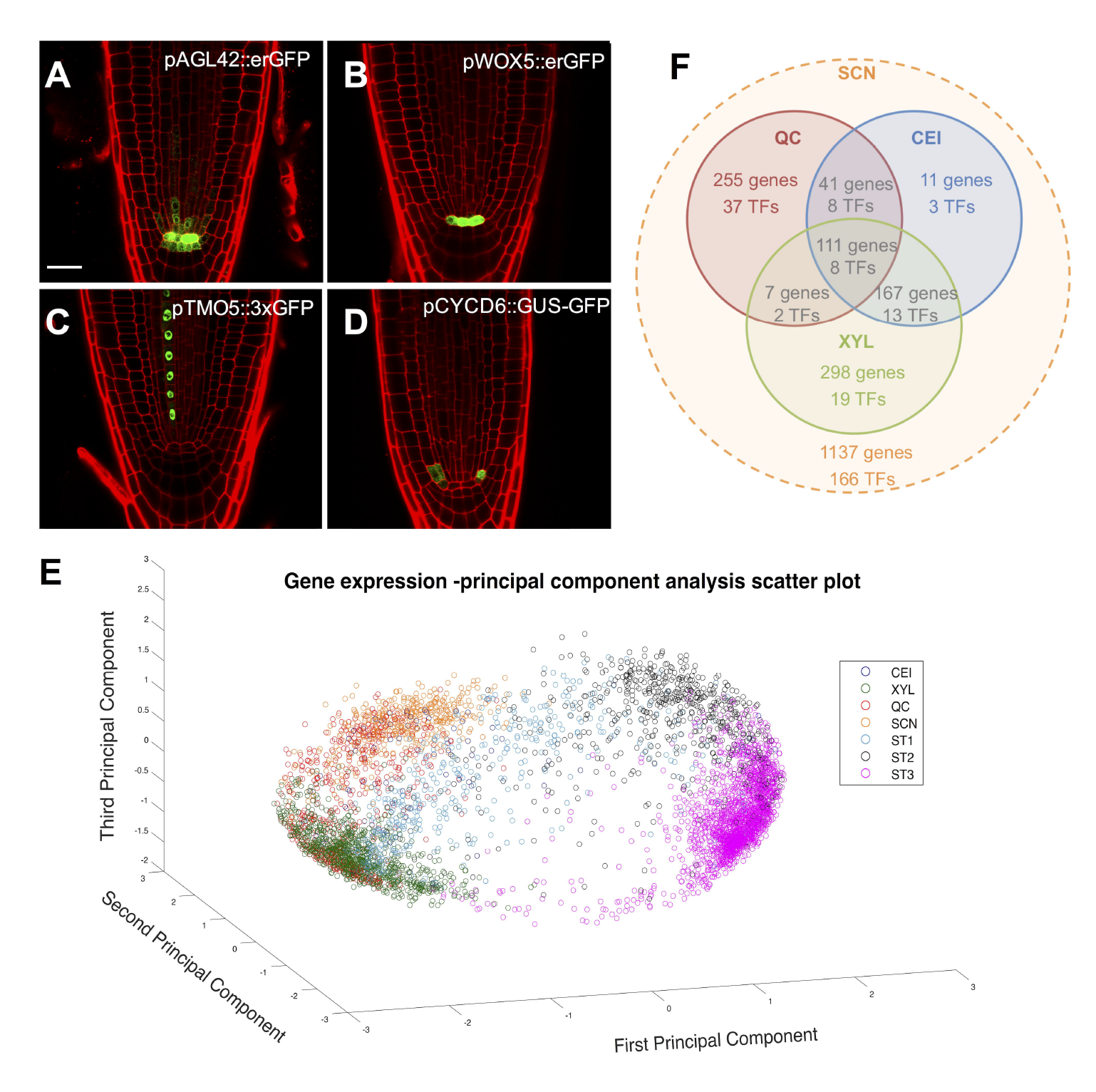
Markers and genes differentially expressed in the root stem cell niche. **A-D** Marker lines for the SCN (A), QC (B), xylem (C), and CEI (D) cells. The SCN encompasses several stem cell populations, which include, among others, xylem, CEI, and QC cells. **E** PCA of the genes enriched in each cell or developmental zone. Genes for the PCA are represented by their expression patterns across cell-type and root zone. Circles represent genes, and colors indicate the cell or zone of enrichment of a particular gene. **F** Venn diagram of the number of genes and TFs identified in each stem cell type. Of the 1625 genes enriched in the stem cells, 402 were found to be enriched in the SCN and in one or more of the cell types (not shown in the diagram for clarity, see Dataset S01).

### *In silico* and *in vivo* validations of GENIST

To predict how the 201 TFs cooperate to regulate the stem cells, we developed a computational pipeline (GENIST) aimed at predicting gene interactions from a combination of spatial and time-series gene expression data. GENIST was developed to predict and prioritize interactions (edges) and key genes (nodes) by integrating two consecutive computational strategies: a clustering step and a Bayesian network inference step (Supplemental information, Fig. S1), which use spatial and temporal datasets, respectively. Since GENIST’s inference step and its integration with clustering had not been previously tested, we performed a modular validation by testing first the DBN inference step, and next, the incorporation of clustering into the algorithm. We tested GENIST’s inference step with *in silico* time-series datasets (DREAM 4 challenge 2 (23–25)) and found that our algorithm outperformed previously published methods (ebdbnet (26), ScanBMA (27), ARACNE (28), CLR (29), MRNET (30), LASSO (31,32)) in terms of precision and Area under the Precision Recall Curve (AUPRC) (Supplemental information, Tables S2-S3). We next tested whether GENIST could recover known root networks by inferring a phloem (Fig. S2A-E), a CEI (Fig. S3A-D), and a XYL network (Fig. S3E-H) (See supplemental information for details). Our inferred networks (Fig. S2E, Fig. S3B, F) outperformed previously published methods (ARACNE (28), CLR (29)) (Supplemental information) and showed that GENIST could be used to infer root GRNs. Further, we validated the integration of the clustering and inference steps by testing the performance of both, GENIST’s inference step alone, and the integration of the clustering and the inference steps, to infer regulations among genes expressed across distinct cell types. For this, we applied GENIST to a dataset that compressed the previously tested phloem, CEI, and XYL networks together. We found that the inference step alone resulted in a low inference precision (Precision = 0.071) that was improved when GENIST was applied with both of its steps (Precision = 0.5556). This validation confirmed that GENIST can be used to infer networks of genes expressed across different cell types. In addition, this validation indicated that the use of clustering prior the inference step boosts the performance of the algorithm when the network genes are not spatially co-expressed (Supplemental information, Fig. S2F). Furthermore, we showed that clustering improves the algorithm computational complexity (Table S4).

To increase our confidence in our GRN inference pipeline, we performed a validation with our stem cell dataset. Specifically, since several datasets involving XYL-enriched genes, such as ChIP-chip data on SHR (9), were available, we applied GENIST to identify regulatory interactions among the TFs found in the XYL (Fig. S4A), and in turn, downstream of SHR. Among the predicted SHR targets, we found that *AT4G24060* and *GATA5* were directly bound by SHR (Fig. S4B) (9). We also found that *PHB* was an indirect target of SHR (33), and as shown in a time course experiment (9), *INDOLE-3-ACETIC ACID INDUCIBLE 1 (IAA1)* was also downstream of SHR (8) (34). Unfortunately, we were not able to test *NAC076* and *ACETIC ACID INDUCIBLE 29 (IAA29)* since the Agilent chip did not have probes covering the promoter region of such genes (9). Alternatively, we used the expression profiles obtained from *shr* mutant roots (33) and obtained information on SHR regulation sign (activation/repression) for *NAC076, IAA29,* as well as the *AT4G24060* and *GATA5* direct targets (Fig. S4B). Overall, our validation indicated that the capacity of GENIST to integrate spatiotemporal datasets is key for inferring GRNs in organisms where transcriptional datasets of diverse characteristics are available.

### GENIST identifies PAN as a root stem cell regulator

We applied GENIST to our stem cell dataset to infer GRNs and to help us identify additional stem cell regulators. Specifically, we inferred the network among the genes enriched in the QC (Fig. 2A) leveraging our spatial dataset and the transcriptional profiles of the 12 developmental time zones of the root. We then investigated the inferred QC-network to assess its main nodes. Our QC marker, *WOX5,* was not present in the network, since it was not contained in the Affymetrix ATH1 GeneChip used to obtain the expression profile of the stem cells. However, other known stem cell regulators, such as *NO TRANSMITTING TRACT (NTT)* (35), *MYB56 (BRAVO)* (36), *and AIL6* (2) (Dataset S02), were found among the main nodes. Since the main hub, *NTT,* together with *WIP DOMAIN PROTEIN 4 and 5* (*WIP4* and *WIP5),* were recently shown to be essential for root development (35), we studied the *NTT* subnetwork to investigate its downstream hubs. We identified *PAN* as one of the main hubs in the *NTT* subnetwork. Since PAN was previously shown to be involved in a feed-forward loop with *AGAMOUS (AG)* and *WUSCHEL (WUS)* in the shoot apical meristem (37,38) we hypothesized that *PAN* could have a role in regulating the root stem cells.

**Figure 2.**
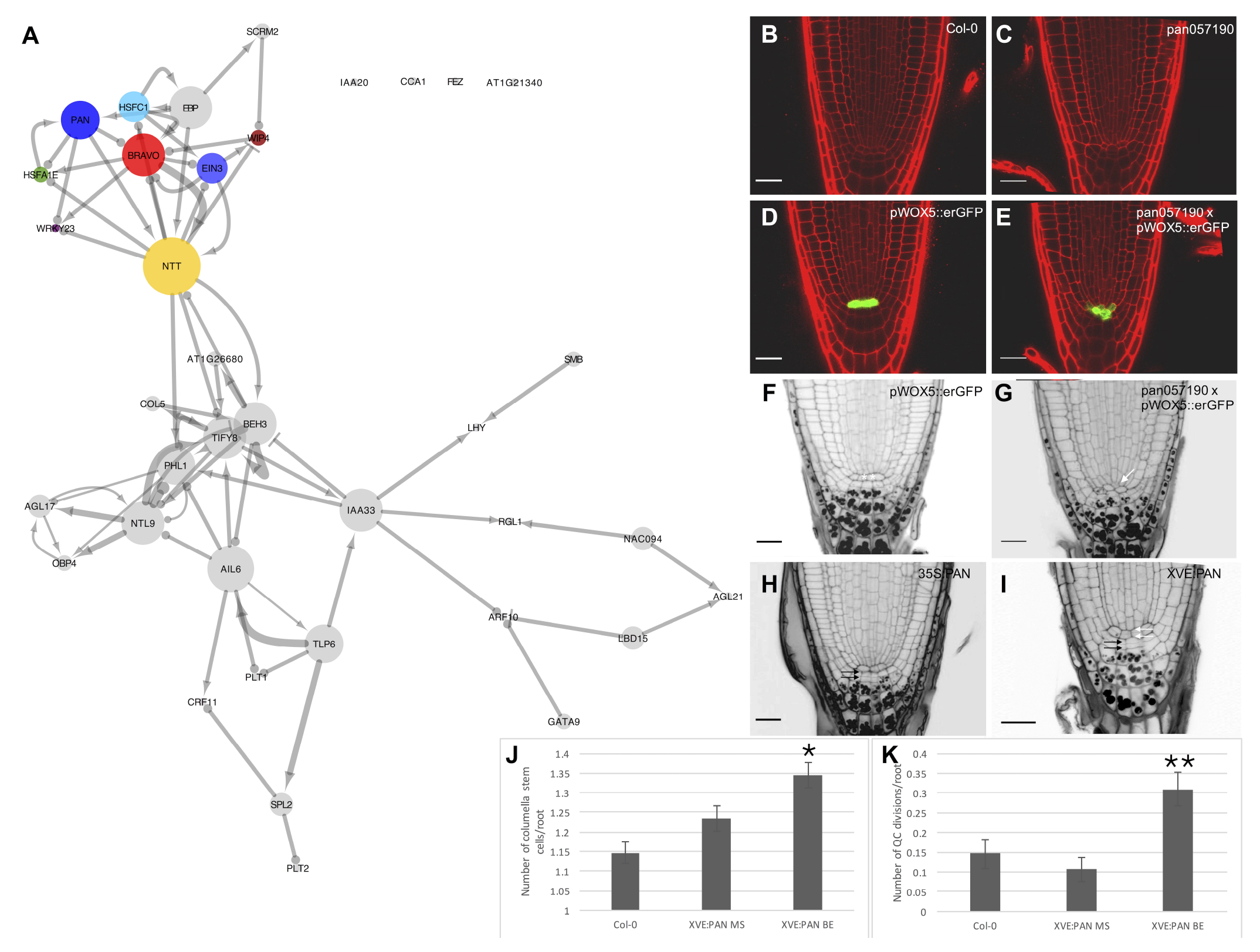
QC TF network. **A** Network among the QC enriched TFs. Node sizes indicate importance of the nodes in terms of the number of TFs that they regulate. Color-coded nodes represent genes downstream of PAN that were used for the mathematical model and experimental confirmations. **B-I** Confocal images of 5-day old Arabidopsis roots **B** Col-0 wild type root **C** *pan057190* root showing a disorganized SCN D *pWOX5:erGFP* root **E** *pWOX5:erGFP;pan057190* root showing changes in QC marker expression **F-I** Confocal images with mPS-PI staining for starch granules **F** *pWOX5:erGFP* root **G** *pWOX5:erGFP;pan057190* root showing differentiated columella stem cells **H** *35S:PAN* root showing extra columella stem cell layers **I** *XVE:PAN* root showing QC divisions and extra columella stem cell layers **J** Quantification of the number of columella stem cell layers in the different lines K Quantification of the number of roots showing QC divisions. Star (*) represents a significant statistical difference (p<0.05, Wilcoxon rank sum test) between *XVE:PAN* upon ß-estradiol treatment (BE) and Col-0 WT control. Double star (**) represents a significant statistical difference (p<0.05, Wilcoxon rank sum test) between *XVE:PAN* upon ß-estradiol treatment (BE) and *XVE:PAN* control treatment (MS). **J-K** Number of roots examined: WT Col-0, n = 41; *XVE:PAN* MS, n = 47; *XVE:PAN* BE, n = 55. White arrows indicate QC cells and black arrows indicate columella stem cells.

We compared the root SCN of three previously characterized *pan* mutant alleles (37–39), a ß-estradiol-inducible *XVE:PAN* transgenic line (40), and a constitutive *p35S:PAN* line (37) (Fig. 2C,E,G-I), with those of control roots (Fig. 2B,D,F). We found that the three *pan* mutant alleles showed a disorganized QC with no discernable columella stem cells with a penetrance similar to what was observed in floral development (12% of *pan057190 (n* = 145), 4% of *pan031380* (*n* = 48), and 10% of *pan247* (*n* = 47)) (Fig. 2C,E,G and Fig. S5A-C). In line with these observations, the *35S:PAN* line and 34.5% of the *XVE:PAN* transgenic plants (*n* = 55) showed additional columella stem cell layers compared to WT Col-0 (*n* = 41) (Fig. 2J). *XVE:PAN* transgenic plants also showed a significantly increased number of QC divisions, with 23.6% of them presenting QC divisions compared to 8.5% in the untreated *XVE:PAN* (p < 0.05) (Fig. 2K). In agreement with the observed phenotypes, the transcriptional fusion of the *pPAN:GFP* line was detected in the QC and adjacent cells (41) (Fig. S5E). Since WOX5 has been shown to be involved in columella and QC function (42), we investigated whether PAN and WOX5 could function in the same pathway. For this, we analyzed the expression of *PAN* and *WOX5* in *wox5* and *pan* mutant backgrounds, respectively (Fig. S6). We found that the expression of *WOX5* was significantly reduced (p = 0.03) in the *pan* mutant background (Fig. S6A), while no change in *PAN* expression was observed in the *wox5* mutant (4). Together, our observations suggest that PAN, upstream of WOX5, is involved in QC function, particularly promoting QC divisions and in turn affecting columella stem cell maintenance.

To gain insight into the dynamics of *PAN* regulation, we constructed a mathematical model of PAN and its predicted downstream targets (Fig. 3A) (see Supplement). Consequently, to model the expression of the 8 TFs over time, we used eight ordinary differential equations (odes) based on Hill equations. We then simulated the fitted system of ODEs with a controlled increase of *PAN* until all the TFs reached their steady states (WT simulation, Fig. 3B) and analyzed how these steady states changed in the absence of *PAN (pan* simulation, Fig. 3C). We found that the steady states of all targets were decreased (converging to zero) in the *pan* mutant with respect to the wild-type simulation, indicating that PAN activates all its downstream direct and indirect targets. In the WT simulation, all targets reached steady states by day one with subtle changes of expression during the transients (time length until expression values reach their steady states). On the contrary, the *pan* mutant simulation showed that *EIN3* and *WIP4* presented high expression values during the transients and reached steady states at later stages (days 3 and 4, respectively). These delayed responses and initial activations of *EIN3* and *WIP4* suggest that, as predicted, these genes could be indirectly affected by *PAN.* Further, the dynamics of our simulations support that BRAVO, NTT, and WIP4 may be connected through feedback loops. During the transient of the mutant simulation, *NTT* and *BRAVO* show an exponential decay, which is consistent with the prediction that they activate each other in the absence of *PAN.* However, their steady states are not immediately reached, since they are activated by *WIP4* and *EIN3.* Conversely, *WIP4,* which is repressed by a decaying *NTT,* shows high levels of expression. Overall, our model predictions suggest that PAN regulates QC function through the activation of *BRAVO*, *NTT,* and *WIP4.* In turn, since these factors are connected through feedback loops, our model suggests the two pathways where these factors function could be interconnected.

**Figure 3.**
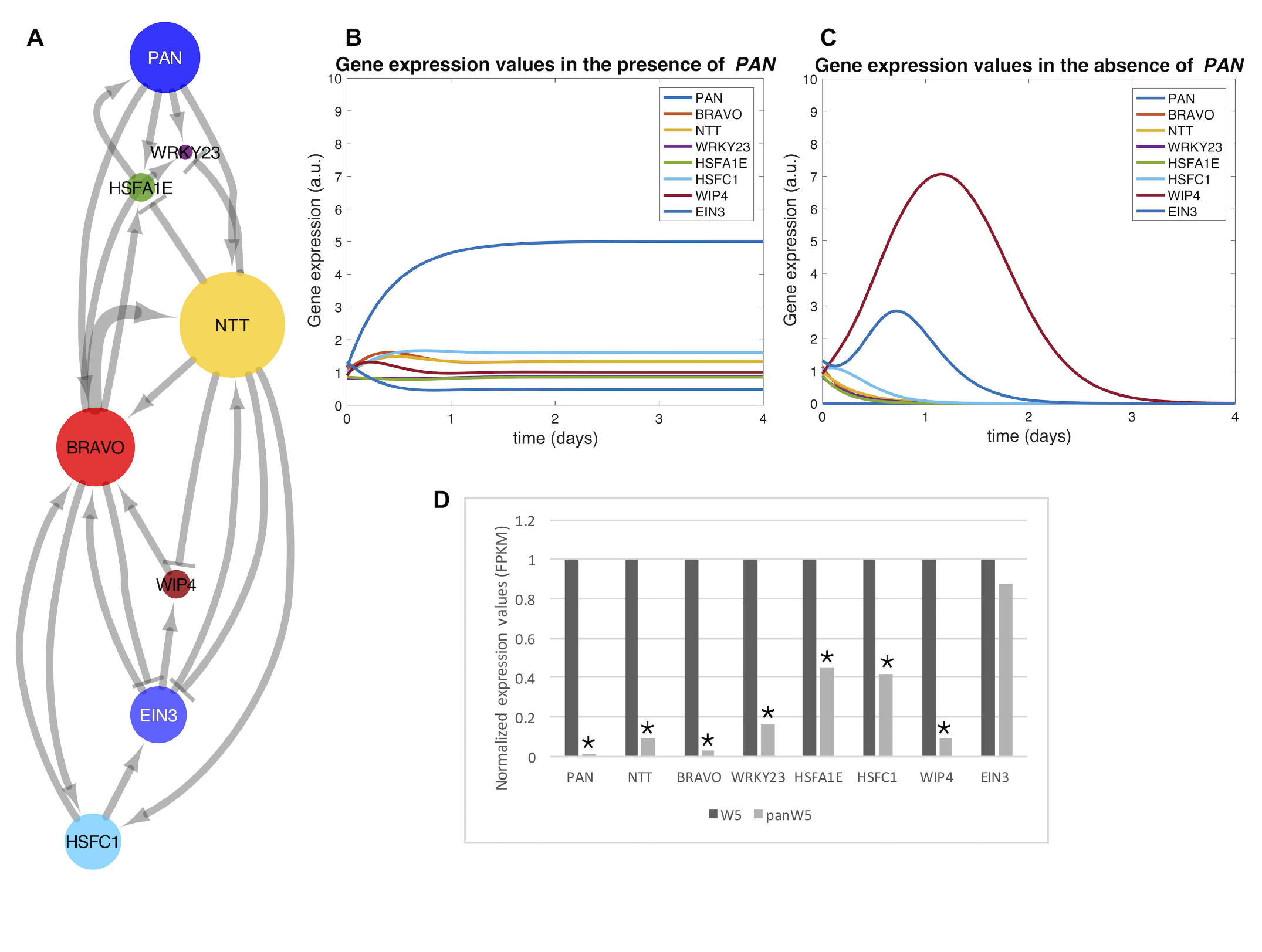
PAN subnetwork in the QC. **A** Optimal configuration of the subnetwork of *PAN* and its downstream targets. **B-C** Resulting expression values of *PAN* and its downstream targets, over time (4 days), after simulating the optimal configuration of the model. **B** Model simulated with the fitted equation parameters **C** Model simulated with the *PAN* associated parameters set to zero to simulate a *pan* mutant situation **D** Normalized expression values of *PAN* and its predicted downstream targets in Col-0 wild-type and in a *pan* mutant. Star (*) represents statistically significant changes of expression between the mutant and the wild type (q < 0.05).

### PAN controls stem cell regulation through an interconnected cellular network

To validate the inferred downstream genes and identify additional factors regulated by PAN, we transcriptionally profiled cells marked by the *pWOX5::erGFP* in a WT and *pan057190* mutant (Methods) (Fig. 2E). We specifically used *pan057190,* as this line showed the largest decrease in relative fold change expression in *PAN* mRNA (Fig. S5D). We found that all the inferred downstream genes of *PAN,* namely *NTT, WIP4, BRAVO, WRKY23, HSFA1E,* and *HSFC1,* with the exception of the indirect gene *EIN3,* showed a decrease in expression in the mutant (q < 0.05 & FC > 2) (Fig. 3D). Thus, the transcriptional profiles in the *pan* mutant, together with the GRN inference and the mathematical model, suggest that *PAN* regulates genes important for QC function. Furthermore, we identified 3397 genes differentially expressed in *pan057190;pWOX5::erGFP* when compared to *pWOX5::erGFP* (Dataset S03), suggesting that *PAN* function affects multiple factors and pathways and might be an important regulator in the stem cells. Accordingly, we found that 75 of the stem cell enriched factors, including key stem cell regulators such as *WOX5, PLT1-2, AIL6, BBM, SHR, SCR, MGP, PHB, FEZ, SOMBRERO (SMB)* (Fig. S6C) (Dataset S03), were differentially down-regulated in the *pan* mutant (q < 0.05 & FC > 2). Thus, to investigate the regulatory effect of *PAN* in the root stem cell niche, we applied our computational pipeline and identified interactions among the 201 stem cell enriched TFs (Fig. 4). We then studied the stem cell network topology to understand which sub-networks could be regulated by *PAN* and whether there is redundancy within these sub-networks. We found that the distribution of clustering coefficients of each node, which measures its connectedness, followed a decreasing power law (Fig. S7A), suggesting that our stem cell network contains highly connected sub-networks (cliques) that confer high levels of redundancy (43–45). By studying these cliques, we found that our network captured stem-cell-specific sub-networks of genes (groups of genes enriched in the same stem cell population) and functional sub-networks of genes (groups of genes that function in the same or related pathways) (Dataset S04). We observed that most of the factors differentially expressed in the *pan* mutant were found in two subnetworks containing QC factors, one subnetwork with embryo and seedling development factors (see Dataset S04), and one subnetwork containing lateral root cap factors. This suggests that *PAN* function may not extend outside of the QC and columella stem cells. Taken together with the *pan* and *XVE:PAN* phenotypes, these results indicate that *PAN* could be involved in QC function and suggests that *PAN* activates a cascade of interconnected factors (downregulated in the *pan* mutant) that control these functions.

**Figure 4.**
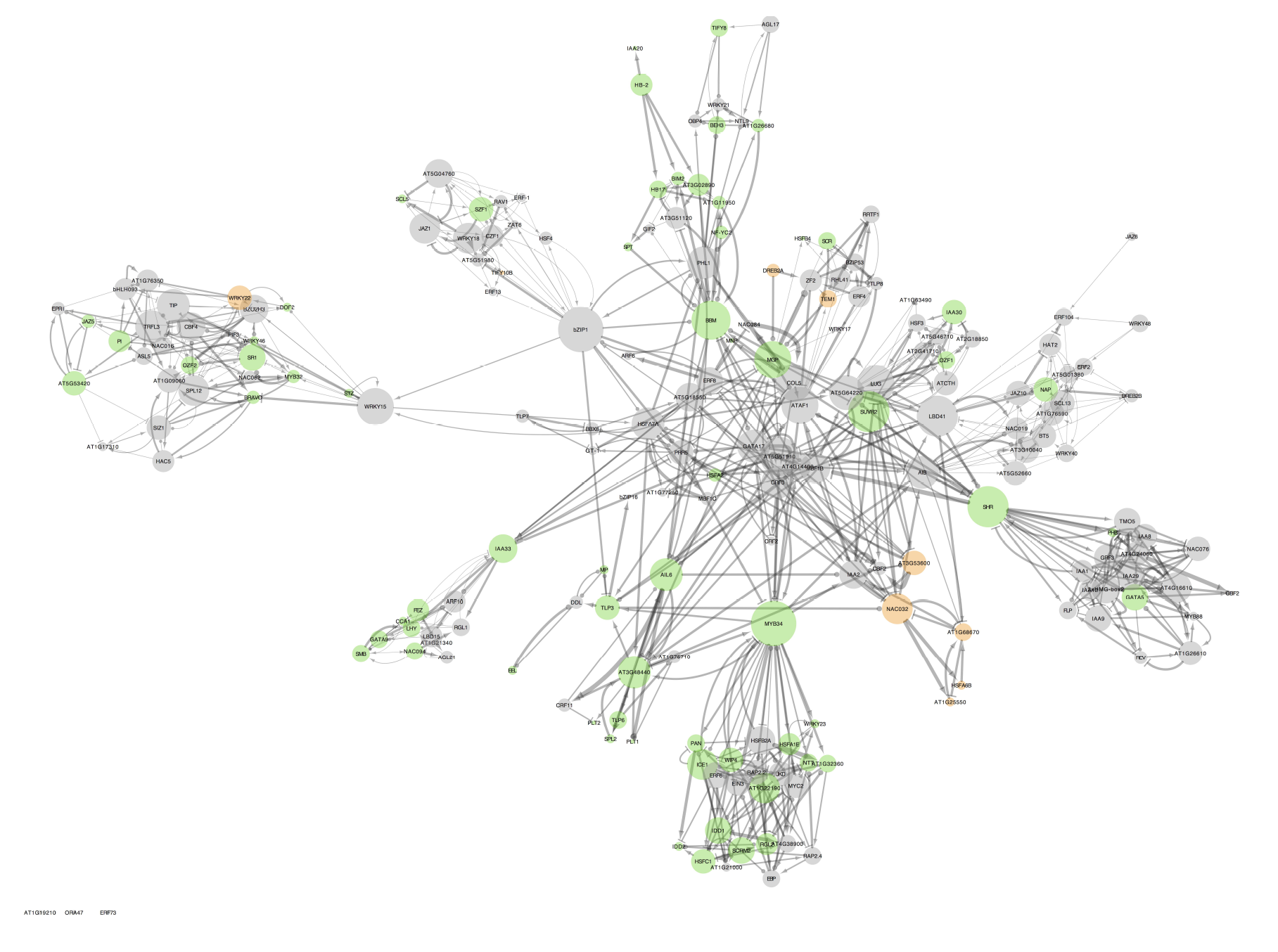
TFRN of the 201 TFs enriched in the SCN. Clusters of nodes indicate groups of TFs functionally related or functioning in the same cell type. Node sizes indicate importance of the nodes in terms of the number of TFs that they regulate. The highly connected groups of genes or sub-networks correspond to the DBN inferred for each cluster. Green (orange) nodes represent factors that are differentially down-regulated (up-regulated) in the *pan* mutant with respect to Col-0 wild-type.

## Discussion

In this work, we took an unbiased approach to identify the transcriptional signature responsible for regulating Arabidopsis root stem cells. We showed that known stem cell regulators do not appear to be differentially expressed in the stem cells with respect to the meristematic zone. This suggests that “stemness” might be a property that gradually decreases as cells move further from the QC. This findings allowed us to use the stem cell gene expression along the root axis to infer DBNs. In turn, this allowed us to developed GENIST and showed how its capacity to integrate spatial (stem cell datasets) and temporal (longitudinal root datasets) transcriptional datasets can boost the performance of inference algorithms and led to the generation of inferred stem cell networks.

Next, we used the topology of our networks to identify key regulators of the root stem cell niche. Specifically, we found PAN, which is involved in the shoot apical meristem with AG and WOX5 homolog, WUS, to be a potential regulatory node of the QC upstream of *WOX5.* Our observations suggest that, similar to WOX5 (42), PAN controls columella stem cell maintenance and QC division. However, contrary to WOX5, which restricts cell division in the QC (42), our results indicate that PAN induces QC division. This suggests that a balance between these two factors is needed to maintain the quiescence of the QC, or alternatively, that PAN and WOX5 redundantly control QC division and columella maintenance through different networks and in turn PAN regulates genes independently from WOX5. Accordingly, our mathematical model allowed us to understand that PAN activates *BRAVO, WIP4* and *NTT,* suggesting it controls the QC and columella stem cells through these factors. Moreover, the feedback loops connecting these factors in our model suggest that the pathways where these factors function could be interconnected. In addition to the predicted downstream targets of PAN, 75 TFs (including *WOX5*), whose expression is enriched in the stem cells, were found to be differentially expressed in the *pan* mutant, suggesting that PAN activates a cascade of factors involved in stem cell regulation.

Our combined approach of cell-type whole genome expression analysis, development of the GENIST computational pipeline, and mathematical modeling led to successful findings of factors that could play important roles in stem cell regulation and, in particular, QC function. The finding of PAN as a potential key regulator of the QC function can guide future work to further understand regulatory mechanisms underlying stem cell regulation. We foresee that although much work remains to be completed to determine the role of PAN regulating the QC function, our initial findings will be key for understanding the networks underlying stem cell regulation. Overall, our results highlight the power of the inference tools for exploring gene candidates of interest. In addition to the findings that were shown in this work, our networks have the potential to guide future stem cell research in the Arabidopsis root by identifying genes for further experimental validation. We anticipate that our experimental and computational approach can be applied to solve similar problems in a diverse number of systems, which can result in unsupervised predictions of gene functions and gene candidates.

## Methods

### Plant Material and Growth Conditions

*A. thaliana* seeds were plated and grown in a vertical position at 22°C in long-day conditions (16 hour light/8 hour dark cycle) on 1X MS (Murashige and Skoog) medium supplemented with sucrose (1% sucrose total). T-DNA insertion lines for *PAN* (At1g68640) were used and homozygous lines were confirmed using the primers listed in Table S5. The three *pan* mutant alleles used were *pan057190* (SALK_057190(46)), *pan031380* (SALK_031380(47)), and *pan247* (SAIL_247(48)), with T-DNA insertions in the 3rd intron, 5’UTR, and the 7th intron, respectively). The *XVE:PAN* transgenic homozygous line under the control of a ß-estradiol inducible promoter was obtained from the TRANSPLANTA collection (40).

### Stem cells transcriptional profile

The following GFP marker lines were used to sort the following cell types: QC (*pWOX5:erGFP*(4)), CEI (*pCYCD6::GUS-GFP*(9)), XYL (*pTMO5:3xGFP*(49)), and SCN (*pAGL42:GFP*(10)). Additionally, *pan057190* mutant plants were crossed with pWOX5:erGFP (4). Approximately 250-500mg of seeds per marker line were sterilized using 50% bleach and 100% ethanol. Seeds were imbibed and stratified for 2 days at 4 °C. Afterwords, the seeds were plated on 1X MS agar (+1% sucrose) on top of Nitex mesh. Plants were grown as described above for 5 days. After 5 days, roots were cut and protoplasts were extracted(50). GFP-marked protoplasts were isolated using the Diva cell sorter or the Beckman Coulter MoFlo XDP Cell sorter. Between 1000-2500 protoplasts were collected for each marker line.

RNA was extracted from *pWOX5:erGFP*(4), *pCYCD6::GUS-GFP*(9), *pTMO5:3xGFP*(49), and *pAGL42:GFP*(10) protoplasts using the Qiagen RNeasy Micro Kit. RNA probes were labeled using the AffyGeneChip3’IVTExpressKit (http://www.affymetrix.com/estore/catalog/131549/AFFY/HT+3%27+IVT+Express+Kit#1_1) and hybridized on the Affymetrix ATH1 GeneChip. This was repeated in triplicates for each of the marker lines.

RNA was extracted from ~500 WT and *pan057190* protoplasts expressing *pWOX5:erGFP(4)* using the RNeasy Micro Kit (Qiagen). cDNA synthesis and amplification was performed using the SMARTer Low Input RNA Kit for sequencing. Libraries were prepared using the Low Input Library Prep Kit and sequenced using an Illumina HiSeq 2500 sequencing machine, with 100bp single end reads. Adapters and low quality reads were filtered out using fastq-mcf, in ea-utils software (51). A window size of 5bp for quality trimming was used. Reads longer than 30bp with a quality score greater than 30 were kept. Clean reads were mapped against the TAIR 10 reference genome using the TopHat v2.0.13 (52). Transcripts were assembled using Cufflinks (version 2.1.1) (53). This was repeated for four replicates for each line.

### Gene selection

A mixed-model ANOVA was used to determine the mean expression for each gene in each cell or developmental zone across 3 biological replicates(33). We defined enrichment of expression of a gene in one cell or developmental zone as being highly expressed in that cell/zone relative to all other non-overlapping samples (>1.2 fold expression & q<0.0001 or > 2 fold expression)(9,12). Accordingly, the conditions for SCN enrichment selection were enrichment in the SCN (*AGL42*) relative to the two developmental zones (Stage II and Stage III). The conditions for one or more stem-cell-type enrichment selection were enrichment in those stem cell types relative to all other samples, excluding the SCN. The low number of CEI-enriched genes obtained with these conditions resulted from the expansion of our QC marker, *WOX5::erGFP,* into the CEI (Fig. 1A).

To find genes regulated by *PAN,* we used the WT *vs.* the *pan057190* mutant dataset. Cuffmerge (53) was used on the output of Cufflinks to merge all the assembled transcripts and create a single merged transcriptome annotation. Testing for differential expression between the WT and *pan* mutant lines was performed with Cuffdiff (53). Genes were considered to be differentially expressed between both conditions if they met a q value and an expression fold change condition (q<0.05 & > 2 fold change).

### Quantitative RT-PCR

Seeds were plated and grown for 5 days as described above. Root tips (2-3 mm) were collected and sonicated to disrupt the cell roots. RNA was extracted using the Qiagen RNeasy Micro Kit. Each RNA sample was reverse transcribed using the SuperScript 3 First-Strand Synthesis System for RT-PCR (Invitrogen) according to the manufacturer’s instructions. qRT-PCR was performed in duplicates for each RNA sample and 3 biological replicates were tested using SYBR green PCR Master mix (Applied Biosystems). Expression levels were calculated relative to *UBQ10* (At4g05320) using the 2-△△ct method. Primers used were either previously published(38,54), or designed using Primer 3 Plus, and are listed in Table S5.

### Microscopy and phenotypic analyses

Phenotypic analysis of WT, *pan* mutant alleles, *XVE:PAN,* and *35S:PAN* roots 5- to 7- days after germination stained with propidium iodide were performed using confocal laser scanning microscopy (LSM710). Images were taken with ZEN software (ZEISS).

Starch granules and cell walls were stained with the mPSPI method and imaged with a confocal microscope as described by Truernit et al. (55). Whole seedlings were fixed overnight at 4°C. The seedlings were then transferred to 80% ethanol and incubated for 3 min at 80°C. Seedlings were transferred back into fixative and incubated for 1 hour, after which time the plants were rinsed with water and incubated in 1% periodic acid for 40 min. The seedlings were then rinsed with water again and incubated in Schiff reagent with propidium iodide for 5 minutes. The Schiff reagent was removed and seedlings were incubated in 500 uL of chloral hydrate overnight. The seedlings were removed from the chloral hydrate solution and mounted on slide using 75 uL of Hoyer’s solution. The number of observable columella stem cell (CSC) layers and the number of QC divisions after mPS-PI staining were counted for the phenotypic analyses.

Multiple plants were imaged to quantify the specific fluorescent markers. 13% of our *pWOX5:erGFP* marker showed expression specific to the QC cells (n = 23) while 61% showed high expression outside the QC cells and particularly in CEI cells, and 26% showed low expression outside in CEI cells. The expression of *pWOX5:erGFP in pan057190* extended into the CEI cells as well. *TMO5:3xGFP* expression showed a graded profile with high expression closer to the QC that gradually decreases. This graded expression enabled the selection of only the first 4 to 8 cells closer to the QC.

## Acknowledgments

We thank Bob Franks of NCSU for providing the *pan* alleles, and Wolfgang Busch for his help with the ChiP-chip SHR targets validation. Support for this work was provided by the National Science Foundation (NSF CAREER MCB 1453130) to RS; MINECO (BIO2014-57107-R), CONSOLIDER (CSD2007-00057), Junta de Castilla y León (SA093U16) and ERC.KBBE.2012.1.1-01 (EcoSeed-311840) to OL. MGF-E is supported by a JCyL and Fondo Social Europeo grant. Funding to APF is provided by NSF (DGE-1252376). Funding to NMC is provided by NSF (DGE-1252376). Funding to BKM was provided by an EMBO-STF (ASTF4-2012). The data presented in this paper have been deposited in NCBI’s Gene Expression Omnibus and are accessible through GEO Series accession number GSE15876, GSE64253, GSE76710, and GSE97792.

